# ABC_GP_IPM:A Python package for applying GP and ABC to Integral Projection Models, with a Soay sheep case study

**DOI:** 10.64898/2026.01.08.698488

**Authors:** Zhixiao Zhu, Maria D. Christodoulou, David Steinsaltz

## Abstract

The integration of Gaussian Process (GP) models with Approximate Bayesian Computation (ABC) has been explored as a flexible framework for constructing Integral Projection Models (IPMs), enabling non-parametric modelling of demographic relationships and the incorporation of population-level information without explicit likelihoods. However, the practical implementation of this framework – particularly the selection of ABC summary statistics and the execution of ABC-PMC samplers – remains non-trivial and can limit its broader adoption.

To address this gap, we introduce `ABC_GP_IPM`, a `Python` package that provides a streamlined and user-friendly interface for constructing GP- and ABC-based IPMs. We demonstrate the utility and performance of the package through a real-world case study of Soay sheep (*Ovis aries*), illustrating how the software simplifies complex modelling workflows and data management in IPMs.

## 1 Introduction

Population dynamics research frequently confronts the challenge of modeling complex biological systems with inherent variability and uncertainty. Our previous work improves Integral Projection Models (IPMs) by integrating Gaussian Process (GP) models via a custom Approximate Bayesian Computation (ABC) sampler, enhancing their adaptability and the ability to reflect vital rate interactions (Zhu et al., 2025a,b). However, the practical implementation of this innovative approach is non-trivial, requiring considerable computational expertise, which can be a barrier for researchers without a strong programming background.

To address this, we developed a dedicated `Python` (Van Rossum & Drake Jr (1995)) package, `ABC_GP_IPM`, simplifying the execution of our proposed ABC GP IPM. This tool integrate the selection of ABC summary statistics (Zhu et al. (2025b)) and the ABC sampling process (Zhu et al. (2025a)), providing a user-friendly interface that allows researchers to concentrate on their biological inquiries rather than the complexities of computational implementation.

A key feature of our package is its respect for the highly context-specific nature of demographic studies. It allows users to customise functions, like those for IBM simulations, according to the actual life cycle of the species under study. This customisation is not confined to fixed forms but requires adherence to certain rules defined within the package, including specific conventions for naming, input, and output formats. Such structured flexibility ensures that, while the package is adaptable to various ecological contexts, it maintains a consistent framework that facilitates reliable and reproducible research.

The decision to develop this package in `Python` was intentional, primarily to leverage the capabilities of `GPflow` (Matthews et al. (2017)) — an advanced library for building GP models. `GPflow` provides a powerful and flexible framework for specifying and solving GP models with cutting-edge techniques, including integrating of `TensorFlow` (Abadi et al. (2015)) for faster computation and GPU acceleration. By developing in `Python`, our package also benefits from the extensive scientific computing ecosystem available in `Python`, which includes libraries such as `NumPy`, `SciPy`, and `matplotlib` for numerical computations and data visualisation (Harris et al. (2020); Virtanen et al. (2020); Hunter (2007)). Furthermore, Python’s widely use in data science and machine learning makes it accessible to a broader audience, potentially facilitating interdisciplinary research.

Our package has targeted functionality, specifically designed to support the initial stages of demographic modeling (see the workflow in Figure 1). This includes a systematic workflow to select ABC summary statistics (Zhu et al. (2025b)) and the implementation of the tailored ABC sampling process (Zhu et al. (2025a)). While our tool provides robust support for these critical preliminary steps, it deliberately does not include the construction of IPM matrices (Step 5 in Figure 1). This domain of IPMs is already well-serviced by a number of robust and mature packages, particularly within the `R` (R Core Team (2016)), for example, `IPMpack` (Metcalf et al. (2013), `ipmr` (Levin et al. (2021)). Recognising this, we have chosen not to replicate these functionalities but instead focus on enhancing the efficiency of the stages leading up to IPM construction. For researchers needing to construct IPM matrices, our case studies includes `Python` code examples. For those wishing to utilise existing R-based IPM functions or tools directly, we recommend exploring well-established interoperability tools like `rpy2` (Gautier (2023)). These tools support executing R code within `Python`, offering a easy integration for users who wish to combine the strengths of both programming environments.

**Figure 1:**
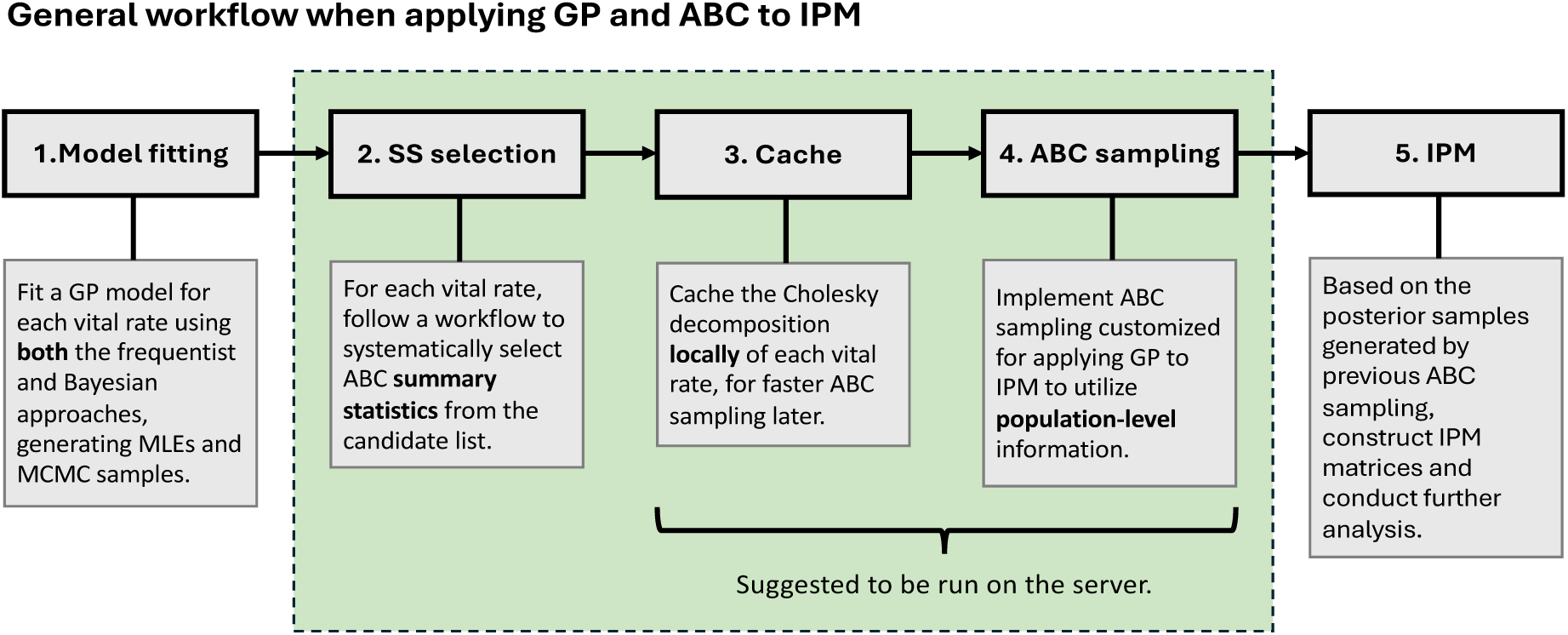
A general workflow when applying GP and ABC to IPM. The processes within the green box have been encoded in our package.

This report starts by an introduction to the foundational background of our study, including the use of object-oriented programming in `Python` (Section 2) and details about the study species (Section 3). We then demonstrate the utility and performance of our package through a case study involving real-world biological data (Section 4), showcasing how our tool simplifies complex modeling processes and data management in IPMs. The results analysis and discussion for the case study are presented in Section 5 and 6, followed by a broader discussion of the package in Section 7.

## 2 Object-oriented programming in Python

The `ABC_GP_IPM` package is implemented following the object-oriented programming (OOP) paradigm with `Python`, which improves code readability, reusability and ease of maintenance. In OOP, a class serves as a blueprint for creating objects, defining their behaviour through methods and setting their initial attributes. An instance refers to an actual object created from the class. For illustrative purposes, consider the initialisation process in `R`:

`human <- list(name = “Marc”, age = 29);`

compare to Python:

`human = Human(name=“Marc”, age=29).`

In this Python example, ‘Human’ represents the class (blueprint), and ‘human’ is the instance (object) constructed with defined attributes ‘name’ and ‘age’.

Python classes encapsulate both data (attributes) and functionalities (methods). For example, the ‘`human`’ instance can perform actions like ‘`eat`’ or ‘`sleep`’ defined within the ‘`Human`’ class, without needing to externally pass attributes like ‘`name`’ and ‘`age`’ each time when these methods are invoked. These attributes are bound to the instance and consistently accessible. This encapsulation allows each instance to operate independently with its own data, promoting a clear and organised code structure.

Additionally, OOP ensures that methods are specific to their relevant classes. For exam-ple, both the ‘`Human`’ and ‘`Sheep`’ classes might have methods for behaviours like ‘`eat`’ or ‘`sleep`’, but each class implements these actions according to its own context. This ensures that methods are relevant and specific to each class.

In the `ABC_GP_IPM` package, we define three classes to handle different use cases, organ-ised hierarchically. For example, the ‘`Human`’ class could be a parent class, with ‘`Female`’ and ‘`Male`’ as sub-classes that inherit common behaviours (methods and attributes). This hierarchy (also called inheritance) allows us to create specific instances of ‘`Female`’ or ‘`Male`’ without duplicating code, making the overall structure more maintainable and efficient.

## 3 Case study: *Ovis aries*

We use the Soay sheep (*Ovis aries*) to demonstrate detailed implementations of ABC_GP_IPM package.

### 3.1 The study site

The case study focuses on the population of Soay sheep residing in Village Bay within the Island of Hirta, part of Scotland’s St.Kilda archipelago. The study species, which are small, horned, and almost deerlike sheep, have thrived in St.Kilda for millennia, with historical records dating back between 3,000 and 4,000 years (Clutton-Brock & Pemberton (2004)). They retain characteristics that are intermediate between modern domestic sheep and their wild ancestors, making them a valuable subject for evolutionary studies (Boyd & Jewell (1974)).

Hirta, the largest island of the St.Kilda archipelago, was inhabited continuously until 1930, after which it was designated primarily as a nature reserve (National Trust for Scotland (2024)). The catchment area of Village Bay, historically the principal settlement area on Hirta, encompasses a range of habitats, including grasslands, cliffs, and sand beach, providing a diverse environment for the Soay sheep (Clutton-Brock et al. (2004b)).

In 1932, 107 Soay sheep were translocated to Hirta, with the population expanding to 1,114 by 1952 (Grubb et al. (1974)). During the study period, approximately 300 of these sheep inhabit the 175-hectare catchment area of Village Bay (Simmonds & Coulson (2015)). The population has been closely monitored since 1985, providing extensive datasets for analysis (Clutton-Brock et al. (1997)). The Soay sheep of Village Bay live in a predator-free environment with minimal interspecific competition and human intervention, closely mirroring the living conditions of their wild ancestors (Catchpole et al. (2000); Simmonds & Coulson (2015)). This unique setting can offer a relatively simplified model for studying natural selection and their population dynamics in a near-pristine environment.

### 3.2 Interplay between Weather, Population density, and Vegetation resilience

The dynamics of the Soay sheep population at the study site are closely linked to food availability, which is influenced by both population density and weather factors (Crawley et al. (2004)).

Weather conditions play a critical role in determining food accessibility and, consequently, the survival rates of these sheep (Catchpole et al., 2000; Clutton-Brock & Coulson, 2002; Ozgul et al., 2009; Coulson et al., 2010). During harsh, wet, and windy winters, wild ungulates experience increased heat loss and reduced foraging opportunities (Forchhammer et al., 1998; Post & Stenseth, 1998; Post & Forchhammer, 2002), resulting in higher mortality rates of the study species (Catchpole et al., 2000; Clutton-Brock & Coulson, 2002). Although spring vegetation biomass can be relatively high following such winters (Crawley et al. (2004)), this does not buffer the population against the direct impacts of severe winter weather on survival (Clutton-Brock et al. (2004a)). Summer conditions, in contrast, are generally favourable due to the abundance of food, resulting in no significant impact on sheep survival during this period (Crawley et al. (2004)).

Population density can exert a substantial influence on food availability through increased intraspecific competition (Clutton-Brock et al. (1997)). During winters, high sheep densities intensify competition for limited food, often resulting in substantial mortality (Clutton-Brock et al. (1996)), particularly among juveniles (Clutton-Brock et al. (2004a)). In contrast, periods of low population density are associated with reduced competition, which in turn promotes relatively better individual health and higher survival rates (Clutton-Brock et al. (2004a)).

The resilience of the vegetation on Hirta is also pivotal in balancing sheep population density and food supply. Winter conditions variably affect vegetation biomass on Hirta, with some years experiencing exceptionally high grazing pressures. Despite these fluctuations, the vegetation consistently demonstrates remarkable resilience — recovering rapidly by the subsequent spring and summer, and by August, achieving biomass levels that are comparable to those observed in typical years (Mysterud (2006); Crawley et al. (2004)). This robust regrowth is critical to support surviving sheep, thereby facilitating their population recovery during the summer months. Although there is significant variability in vegetation biomass in the beginning of spring (March), the biomass at the onset of winter (August) exhibits considerably less variation (Crawley et al. (2004)), indicating a relatively stable food supply during winter. This stability suggests that the primary challenge to winter survival are more likely due to reduced foraging opportunities from harsh weather and intraspecific competition, rather than the food reserves (Jones (2003); Crawley et al. (2004)).

### 3.3 Data collection and the Study dataset

Since 1985, the Soay Sheep Research Project has been using individual tags to track each sheep from birth, through breeding attempts until death, building a robust database of over 8,000 sheep in Village Bay (Team (2024)). Data collection is conducted by the field assistant visiting the island three times per year. Each visit involves comprehensive censuses and searches for deceased sheep, aiming to maintain accurate records of population density and mortality (Clutton-Brock & Pemberton (2004)). The research expeditions are structured around three seasonal focuses:

- **Late Winter and Spring:** Daily searches to monitor winter mortality; newborn lambs are tagged, weighed, and blood sampled shortly after birth.
- **Summer:** Capture 50-70% of the population using netting traps for health assessments. Conduct a comprehensive sheep count to gather data about population density.
- **Autumn:** Monitor the rut, determine the oestrus status of ewes, and capture rams for genetic sampling to confirm paternity and understand reproductive success.

Our study dataset is focused exclusively on *females*, providing detailed individual information spanning from 1987 to 1996. The dataset includes identifiers, age, survival status, body weight for the current and next year, reproductive data (numbers and sizes of *female* newborns), and annual population sizes (which also count *males*) (Coulson (2012)). Our model training process will utilise information from the first seven years (1987 to 1993), while the remaining three will be reserved for model testing.

### 3.4 Data analysis and Model settings

Data entries that were inconsistent with predefined conditions have been systematically removed prior to modeling. For example, each row in the data file is expected to represent a female sheep known to be alive by August census of each year (Coulson (2012)). Entries with missing survival statuses are therefore removed to align with established criteria for living individuals

We further assumed that the observed data for individual sheep each year are representative enough for the unmeasured segments of the population. This assumption enables us to circumvent the complexities associated with an artificially predefined capture process when conducting IBM simulations in ABC sampling. More specifically, for example, unlike deer, Soay sheep show less hiding behaviour to their offsprings; juvenile and yearling females typically follow their mothers closely from birth (Grubb, 1974; Clutton-Brock et al., 2004a). This behavioural trait suggests that the capture of a mother would logically imply the capture of her (newborn) lamb. However, a great number of missing values in newborns for observed mothers can be found in the study dataset. This discrepancy indicates that the capture process may be influenced by factors not fully understood or captured in our dataset. Therefore, by simplifying the capturing process, we facilitate a direct comparison between the simulated populations and the observed datasets, acknowledging the potential nuances and limitations of this setting. In this case with ABC sampling procedure, since we only observed partial information, we applied a bootstrap method to supplement the missing individuals in the initial population structure. For instance, if the population size was 200, but only 130 were observed, we randomly sampled with replacement from these 130 individuals to reach the full population size of 200.

In the study dataset of the Soay sheep population, the occurrence of twin births happens in less than 1.9% of cases. We follow (Simmonds & Coulson, 2015; Coulson, 2012) to exclude it to simplify the modeling process. In this case, we assume that, in the rare event of twins being born, only one female offspring survives to the next census. Our IPMs simplify the complexity of individual variability by representing each sheep solely by its measured size *z*. The population with size *z^′^* at the next time step is described by:

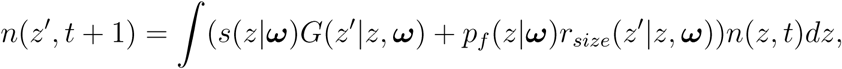

where ***ω*** represents living condition factors influencing vital rates. Specifically, the model describes that a female individual of size *z* has a survival probability *s*(*z|****ω***) and follows a growth trajectory determined by *G*(*z^′^|z,* ***ω***), over an annual cycle ending each August. For reproduction, it starts with considering the probability of recruitment, *p_f_* (*z|****ω***). This is the likelihood that a female of size *z* will become pregnant, then give birth to a *female* newborn, and eventually have that newborn survive to the next year. Conditional on the such recruitment, the female newborn is expected to grow to size *z^′^* with *r_size_*(*z^′^|z,* ***ω***) by the next August, influenced by her mother’s size *z*. Since the data accounts for female newborns only, there is *no* need to set up an additional sex rate for recruitment in our IPM; such a rate is already built into the observed datasets.

Living condition factors encapsulated by ***ω*** include winter weather conditions and initial population densities each August. Given the constraints imposed by the limited seven-year span of our training dataset, strategic modifications about this 2-D vector, ***ω***, were essential. These adjustments address the challenges posed by sparse data points across its continuous variable range. Although such modifications inevitably result in some information loss, they have been carefully crafted to retain critical insights from previous research on the dynamics of the Soay sheep population.

- Winter weather conditions of St.Kilda are largely influenced by the North Atlantic Oscillation (NAO), a climatic phenomenon known to affect many marine and terrestrial ecosystems ((Forchhammer et al., 1998; Belgrano et al., 1999)). The NAO index, reflecting large-scale atmospheric patterns over the North Atlantic (Rogers (1984)), can lead to winters on St.Kilda that are either windy and wet or colder and drier (Clutton-Brock et al. (2004b)). Particularly, stormy winters, indicated by high NAO indices, limit ungulate movement and increase thermal stress in the wild (Forchhammer et al., 1998; Post & Stenseth, 1998; Post & Forchhammer, 2002), which significantly in turn adverse consequences for the sheep population detailed in Clutton-Brock et al. (2004a). To address the marked influence of *high* NAO indices, ***ω*** first consider an averaged NAO index from December to March (Simmonds & Coulson (2015)); then, a floor function is employed to adjust the averaged NAO index, eliminating negative values to focus exclusively on the significant impacts of high positive NAO values on sheep dynamics.
- In addition to the impact of winter weather, our analysis also considers the influence of population density on the population dynamics. While we previously acknowledged that population growth is significantly affected by population density, we observe that this relationship is not linear. Analysis in Clutton-Brock et al. (2004a) reveals a pronounced threshold effect on St.Kilda island: below a population size of approximately 1,100 to 1,200 animals, the population tends to increase by an average of 1.27 per animal per year; however, exceeding this threshold triggers a decline at an average rate of about 0.2 per animal per year. Additionally, a strong positive correlation exists between the populations on St.Kilda island and those in our study site, the Bay area. Building on these insights, we have adopted a threshold value for population density in our models, estimated at *exp*(5.9) according to graphical results presented in Clutton-Brock et al. (2004a). Consequently, in our models, population density is treated as a binary variable: it is coded as 1 if initial population densities in August surpass *exp*(5.9); otherwise, it is coded as 0.

GP models were employed for modelling vital rates (see Appendix A for implementation details).

## 4 Getting start with ABC GP IPM

To enable a easier code reusability and maintenance through the package, the first step is to create an instance of a class that manages data and IPM vital rate models related to the target species.

### 4.1 Initialisation

The initiation of the package involves creating an instance of the ‘`GP_IPM`’ class, encapsu-lating population data and GP models for various vital rates. In the example of Box 1, the constructed instance, ‘`sheep_gp_ipm`’, encapsulates the population data in two key formats: (1) The ‘`popu_data`’ attribute represents the consolidated dataset across all training years, used for model fitting in Step 1 of the workflow in Figure 1. (2) The ‘`POPUdata_dict`’ is the segmented version of ‘`popu_data`’, broken into smaller datasets sorted by consecutive years. It is used later for comparisons with simulated time series datasets. Furthermore, constructing the instance ‘`sheep_gp_ipm`’ requires the fitted GP models for all vital rates, including MLEs (‘`GPmodel_mle`’), Bayesian GPs with priors (‘`GPmodel`’), and the corresponding MCMC samples (‘`MCMC_samples`’).

The package supports the integration of custom functions to adapt to specific data structures and modeling needs. The method ‘`add_XY_compu`’ allows users to add custom functions for extracting data specific to vital rates. For example, as demonstrated in Box 1, the function ‘`XY_m_grw_compu`’ — which processes ‘`popu_data`’ to generate independent and dependent variables for the growth vital rate — has been added to the instance. Similarly, the ‘`add_model_build`’ method allows the user to specify how the GP model for a given vital rate, such as growth, should be initialised via a custom function like ‘`build_new_m_grw`’.

##### Initialisation

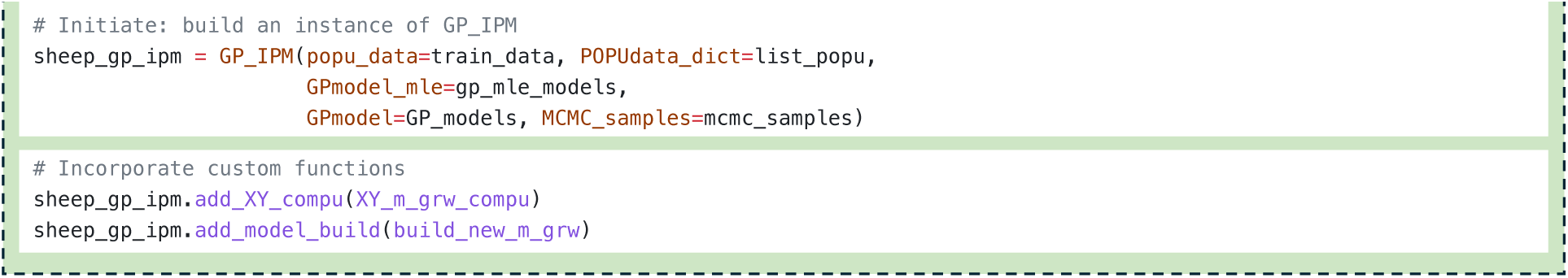

A systematic naming convention is utilised throughout the package to ensure clarity and maintainability. The naming convention employs a prefix “m” followed by the vital rate name to clearly delineate related methods and variables. For example, ‘`GPmodel_mle`’, ‘`GPmodel`’, and ‘`MCMC_samples`’ are `Python` dictionaries with keys such as “m grw”, reflecting models and data pertaining to growth vital rates. This consistent use of naming conventions extends to custom functions, such as ‘`build_new_m_grw`’ and ‘`XY_m_grw_compu`’, ensuring the instance correctly associates each method with the appropriate vital rate. This naming convention will be strictly enforced during implementation, with input validation checks in place to verify adherence.

With the initialisation complete, we make the next three sections correspond to the steps of the workflow shown in Figure 1, providing a clear structure for readers to follow.

### 4.2 SS selection

To select summary statistics for further ABC sampling, the package adopts a systematic workflow based on Negative Log Posterior Probability (NLPP) perturbation (detailed in Zhu et al. (2025b)). This approach selects summary statistics individually for each vital rate.

To determine which summary statistics are more sensitive to shifts within the model, we first need to introduce what aspects of the model are being perturbed and how these pertur-bations are applied. The ‘`Perted_IPM`’ class, a specialised subclass of ‘`GP_IPM`’, is designed to manage these perturbations. In the provided example (see Box 2), ‘`sheep_gp_ipm`’, an instance of ‘`GP_IPM`’, is perturbed specifically for the growth vital rate (‘`m_grw`’) to generate a perturbed IPM object, ‘`sheep_grw_pert`’. Only the first 6% of MCMC samples closest to the optimal NLPP are retained, representing the perturbation results (more details latter).

##### SS selection

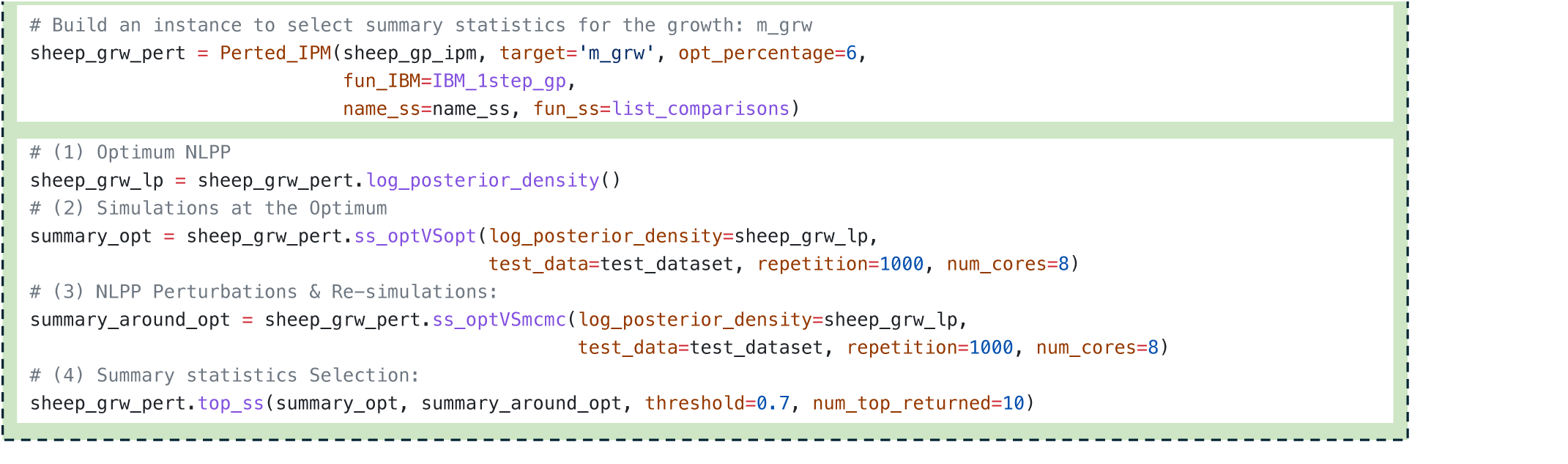

Additionally, to handle diverse scenarios, three user-defined attributes tailored to specific study species are necessary. The function ‘`fun_IBM`’ is used to perform 1-step forward IBM simulations and generate simulated datasets. The ‘`fun_ss`’ function is then employed, computing a list of potential summary statistics and assessing the distance between datasets. These candidate statistics are named as specified by ‘`name_ss`’.

The subsequent steps strictly adhere to the workflow proposed in Zhu et al. (2025b). **(1)** Optimal NLPP: through the constructed perturbed object ‘`sheep_grw_pert`’ with method ‘`log_posterior_density`’, we compute NLPP scores for all MCMC samples to identify which are with or near the optimal NLPP. **(2)** Simulations at the Optimum: we estimate the distance between two datasets simulated under the same hyperparameters associated with the Optimum NLPP through method ‘`ss_optVSopt`’. The argument ‘`test_dataset`’ is used as the initial population structure in a 1-step IBM simulation function to gen-erate ‘`repetition=1000`’ pairs of datasets. This process is set up to use ‘`num_cores=8`’ cores parallel computing to optimise efficiency. **(3)** NLPP Perturbations & Re-simulations: subsequently, though method ‘`ss_optVSmcmc`’, a similar process is followed for the first 6% of MCMC samples closest to the Optimum NLPP (6% was stated when constructing ‘`sheep_grw_pert`’). For each of these samples, we estimate the distance between 1,000 pairs of datasets: one set generated using the hyperparameters at the Optimum NLPP, and the other with hyperparameters from within the 6% closest to the Optimum. **(4)** Summary Statistics Selection: eventually, to evaluate the sensitivity of summary statistics to small deviations from the optimal settings, the area under the receiver operating characteristic curve (AUC) is calculated. An AUC threshold of 0.7 is set to determine sensitivity, and ultimately, the top 10 most responsive summary statistics are reported based on their per-formance by method ‘`top_ss`’. Users can thereby manually select summary statistics based on the returned results.

### 4.3 Cache

To streamline computational efficiency, especially in the context of large-scale simulations in ABC sampling, caching mechanisms are employed. The caching process involves storing the results of computationally intensive tasks, like the Cholesky decomposition needed for GP models, which are frequently used throughout the sampling. With the package, a single line of code activates caching for all vital rates defined within the instance (Box 3). This command stores results in the ‘`Lm`’ directory under the ‘`true_popu`’ parent directory, dynamically adapting to the current script’s execution environment through command ‘`os.getcwd()`’.

##### Cache

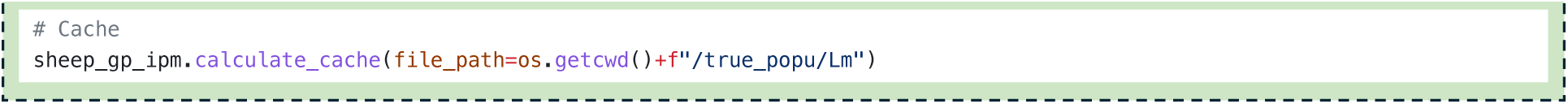

Once the cache are set, the package automatically randomly draws and loads MCMC samples for each vital rate in the background. For each sample, predictive distributions are then calculated in two ways: (1) using the locally stored cache results just computed; (2) recalculating from scratch without the cached data. The package then compares these two sets of results to confirm their consistency, ensuring that the cached computations are accurate and reliable.

Since caching is intended to expedite ABC sampling processes, these caches need to be stored in a location that allows for rapid access by the ABC sampling mechanism. Additionally, due to the large storage demands of caching, it is strongly recommended to execute this program directly on same server used for ABC sampling, rather than execute it locally and then transferring to that server. This way helps to avoid potential delays and resource constraints during ABC sampling..

### 4.4 ABC sampling

We are now ready to proceed to the core phase: ABC sampling. Executing a ABC sampler needs assistance of ‘`ABC_GP_IPM`’ class. At this stage, it is assumed that all necessary preprocessing for ABC have already been completed. Therefore, see Box 4, the attribute function ‘`fun_ss`’ required by ‘`ABC_GP_IPM`’ class is now configured to compute only the list of selected summary statistics. These final listed statistics are named as specified by ‘`name_ss`’. In the sheep example, one summary statistic is chosen for each of the four vital rates.

For IBM simulations, instances of class ‘`ABC_GP_IPM`’ use the ‘`fun_IBM_cache`’ method, which leverages pre-computed cached data for sampling efficiency. To switch from the stan-dard ‘`fun_IBM`’ method to ‘`fun_IBM_cache`’, simply replace the call to ‘`predict_y`’ method with ‘`predict_y_loaded_cache`’ for each of GP models involved in ‘`fun_IBM`’. The method ‘`predict_y_loaded_cache`’ is well-established to speed up sampling, and is readily available for import from our package.

The ‘`fun_IBM_cache`’ conducts IBM simulations using the observed initial independent variables (‘`ini_popu`’). The resultant dataset of this method, representing simulated one-year data, is subsequently processed by the ‘`fun_structure`’ function. This user-defined function is responsible for generating the independent variables needed to initiate the next year’s simulation. Additional parameters for ‘`fun_structure`’ can be supplied through the ‘`fun_structure_kwargs`’ argument. For instance, in our sheep model, ‘`fun_structure`’ also require the observed population density (‘`popsize_list`’) and NAO index (‘`nao_list`’) for the subsequent simulation year (Section 3).

##### ABC sampling

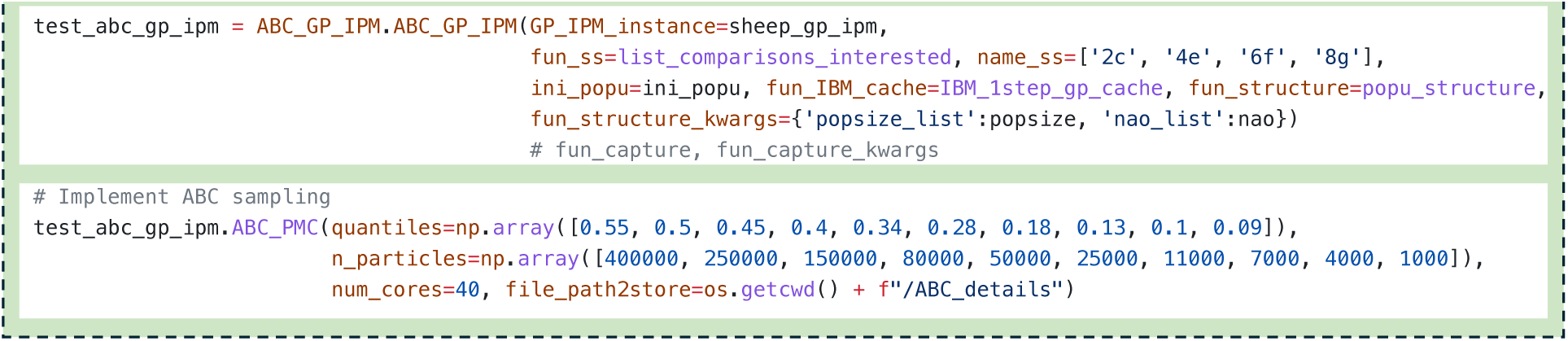

In our Soay sheep case study, as mentioned in Section 3, we assumed that the observed individuals each year are representative enough of the unmeasured population segments. Based on this assumption, we have ignored the capturing process to avoid introducing unnecessary complexity into the simulation. However, users with specific insights or concerns about the capture process can still implement a custom capture function to better reflect their data structure. In this package, the custom capture function can be implemented via the ‘`fun_capture`’ argument. By providing this capture function, the constructed instance is enabled to transform the simulation output of the entire population into a format comparable to the observed datasets during the ABC sampling process. The function should take the simulation output and the simulation year as key inputs, with additional arguments passed through ‘`fun_capture_kwargs`’.

We have finally reached the last step: performing the ABC sampling using the ‘`ABC_PMC`’ method. The algorithm proposed by Zhu et al. (2025a) is all set up and can be easily implemented by specifying the quantiles and the number of particles (‘`n_particles`’). The sampling process in Box 4 is executed with parallel computing of 40 cores. The resulting output files will be stored in the directory specified by the ‘`file_path2store`’ attribute.

### 4.5 Forecasting

The ‘`ABC_PMC`’ method of ‘`ABC_GP_IPM`’ class ultimately results in a set of parameter estimates that represent the approximate posterior distribution, ready for further analysis or predictions. Our package also provide method ‘`simulate`’ for instances belonging to class ‘`ABC_GP_IPM`’, enabling users to easily utilise the results returned from ‘`ABC_PMC`’ to perform population dynamic simulations.

‘`abc_index`’ and ‘`abc_weight`’ are output files of ‘`ABC_PMC`’. The ‘`abc_index`’ is an ar-ray containing indices that reference specific MCMC samples. Each index corresponds to a particular parameter configuration from the estimated ABC posterior distribution. The ‘`abc_weight`’ array holds the associated weights, reflecting the relative importance or likelihood of each sample. During simulation in ‘`simulate`’, the method randomly selects an index from ‘`abc_index`’ based on the probabilities in ‘`abc_weight`’; the corresponding parameter configuration is then used in the simulation of population dynamics.

##### Forecasting

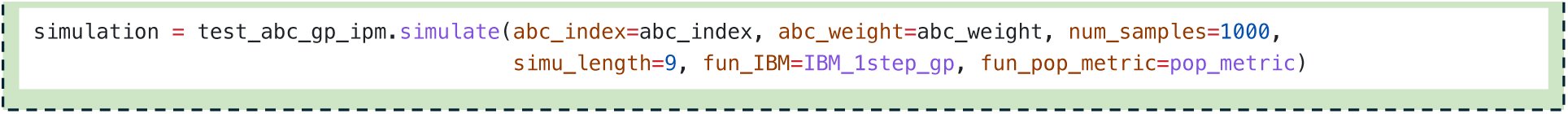

The method ‘`simulate`’ allows the user to specify the number of samples to generate (‘`num_samples`’), the length of the simulation (‘`simu_length`’), and two customisable key functions: ‘`fun_pop_metric`’, which calculates population metrics of interest during the simulation (such as population size or growth rate, etc.), and ‘`fun_IBM`’, which performs user self-defined one-step forward IBM simulation. In our example in Box 5, we are running this method locally, so we used ‘`IBM_1step_gp`’. However, if the method is executed in an environment that allows for rapid access to cached information, it can be also replaced with, for example ‘`IBM_1step_gp_cache`’, to improve efficiency.

The ‘`simulate`’ function may not be an essential part of a general workflow when applying GP and ABC to IPM. However, we include it here not only for users who may find it useful, but also to provide an example for understanding how the results returned from ‘`ABC_PMC`’ should be processed. By inspecting the source code of ‘`simulate`’, users can gain insights on how to handle the output and may even develop further analyses or other types of simulations tailored to their own datasets.

## 5 Case study: Results

We generate ABC samples using the model setup discussed in Section 3 and the code provided in Section 4 via the developed `ABC_GP_IPM` package. These samples are then used to evaluate the ABC GP models’ predictive performance and compared with those from MCMC and GLM methods, as demonstrated in the previous *C. flava* case study.

For the **long-term** forecasting, we initialise the population structure in 1987, using a bootstrap method to account for missing individuals. Predictions are then generated year by year, based on the population simulated from the previous year’s data. For the **1-step** forecasting, we use the observed (incomplete) population structure from the previous year, supplement it with the same bootstrap method, and then make only a single year predictions, keeping the remaining procedures consistent with the approach detailed earlier of *C. flava*. From long-term population forecasting on the Figure 2 left, all models show a general increasing trend in population size over time, with GP and ABC GP closely aligned until 1993, after which ABC GP predicts slightly higher values with larger uncertainty. GLM, while following a similar trend, deviates more from the other two models after 1993, showing wider uncertainty bands. Moreover, all models exhibit increased uncertainty in the later years suggesting the challenges of making long-term predictions over time.

**Figure 2:**
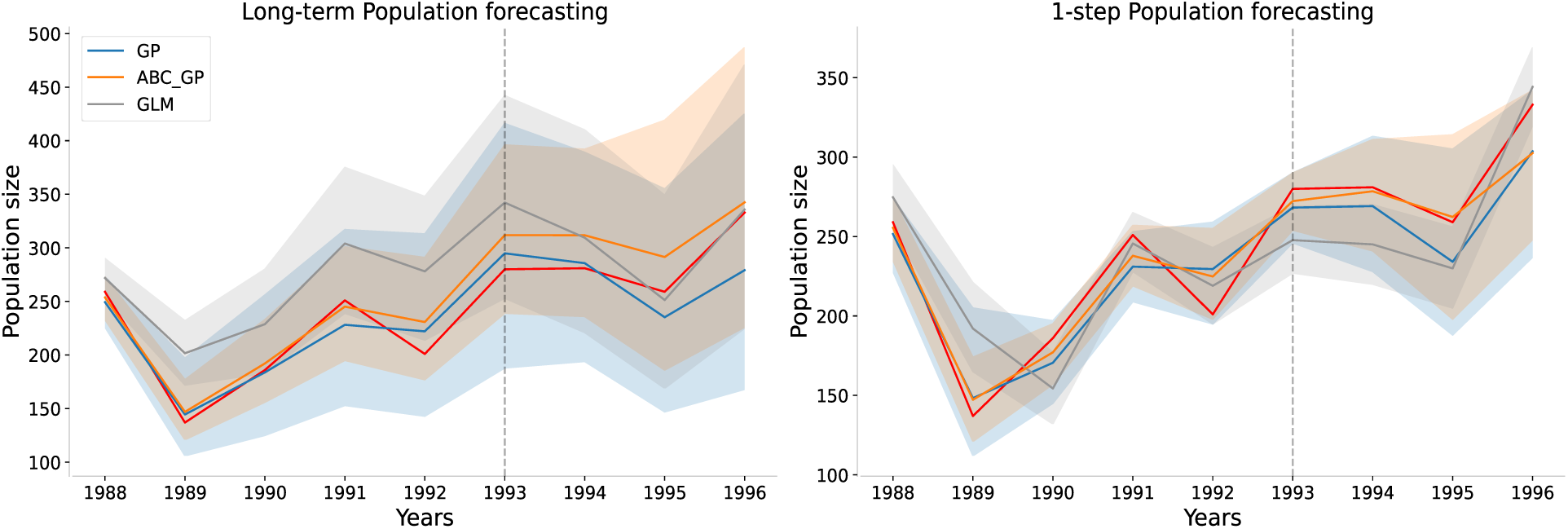
Two types of population size predictions, along with their corresponding 95% P.I. (shaded areas), were generated using GP, ABC GP and GLM samples through 1,000 IBM simulations. The blue predictions were based on MCMC samples, while the orange and grey ones were based on ABC GP and GLM posterior samples, respectively. The point estimations (means) are represented by solid lines in their corresponding colours. The red line represents the true observed population size. The model training process utilised data from 1987 to 1993 (before the grey dashed vertical line). The red line represents the true observed population size.

Compared to long-term forecasting, the 1-step predictions on the right-hand side have narrower uncertainties (as they use more immediate observed data). In 1-step population forecasting, all models display sharper fluctuations, especially in the early years. Here, GP and ABC GP again closely track each other, with ABC GP predicting slightly higher population sizes in the later years. GLM remains the most variable model, with larger deviations and uncertainty.

Let us take a closer look at their point estimations. We use absolute relative differences between the true and predicted population size (Figure 3) to assess each model’s overall performance, without being affected by whether over- or under-estimated. Normalising these differences relative to the true population size ensures that errors are comparable across different population sizes and time points.

**Figure 3:**
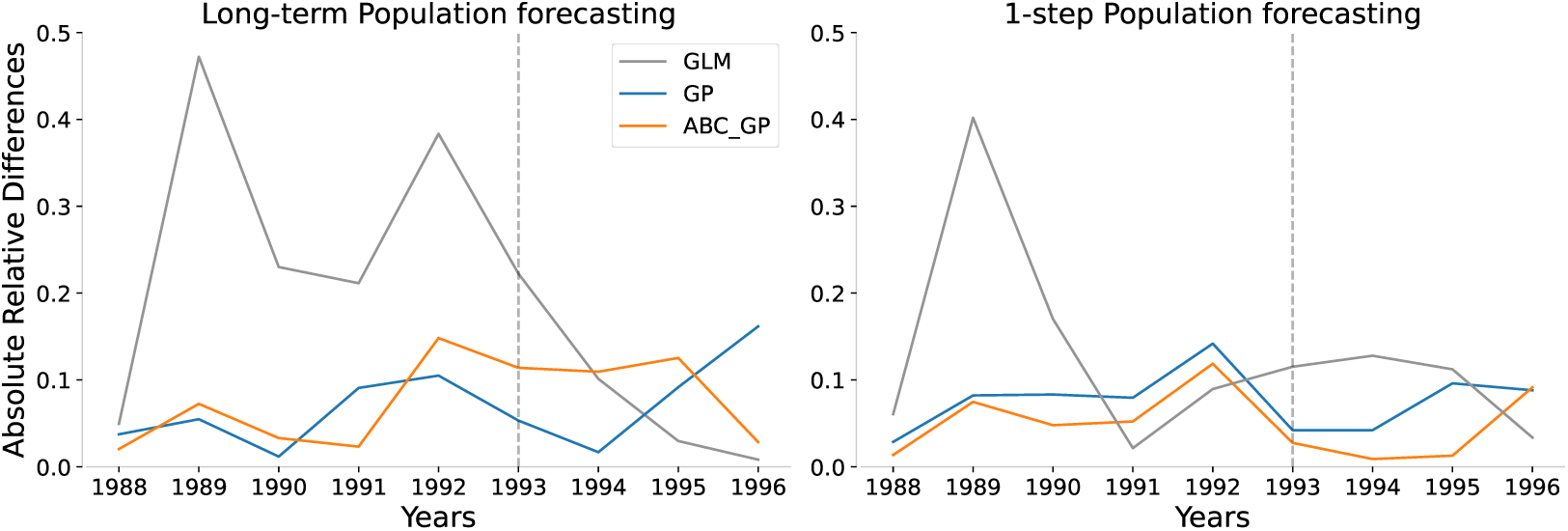
Absolute differences between the observed and predicted population size (means of the predictions) relative to the true observed population size.

Figure 3 left shows the absolute relative differences in long-term forecasting, where both GP and ABC GP models perform well, with the differences staying below 0.2. In contrast, the GLM model shows much larger deviations, peaking around 0.4 in years like 1989 and 1992, indicating relatively less predictive accuracy. The right plot in Figure 3, focused on 1-step forecasting, shows smaller errors across all models compared to long-term forecasting. Both GP and ABC GP maintain strong short-term accuracy, with differences remaining under 0.1 for most years, while GLM again shows more variability.

An interesting point emerges with GLM, particularly in long-term forecasting, where it shows large errors during the training years but surprisingly smaller errors in the test years. However, this may not be a positive sign. The seemingly improved performance in the test period might not reflect GLM’s true predictive power, but rather chance alignment with the test data, raising concerns about the model’s reliability for future predictions.

On the other hand, the GP model performs well during training but shows (slightly) increasing errors in the test years, which might suggest potential over-fitting — the model may struggle to generalise to unseen data.

In contrast, ABC GP stands out for its consistency across both training and test years. While its errors are not always the smallest, they remain stably acceptable across both phases. This stability makes ABC GP, while not perfect, a balanced option for both short-and long-term forecasting.

Figure 4 offers another perspective on the 1-step forecasting results by focusing on the estimated distribution of population growth rates (the one with the similar pattern for long-term is shown in Appendix A). During the training years, the prediction intervals (P.I.) across all models appear fairly comparable in width. However, in the test years, the GP and ABC_GP models tend to have wider P.I. compared to GLM.

**Figure 4:**
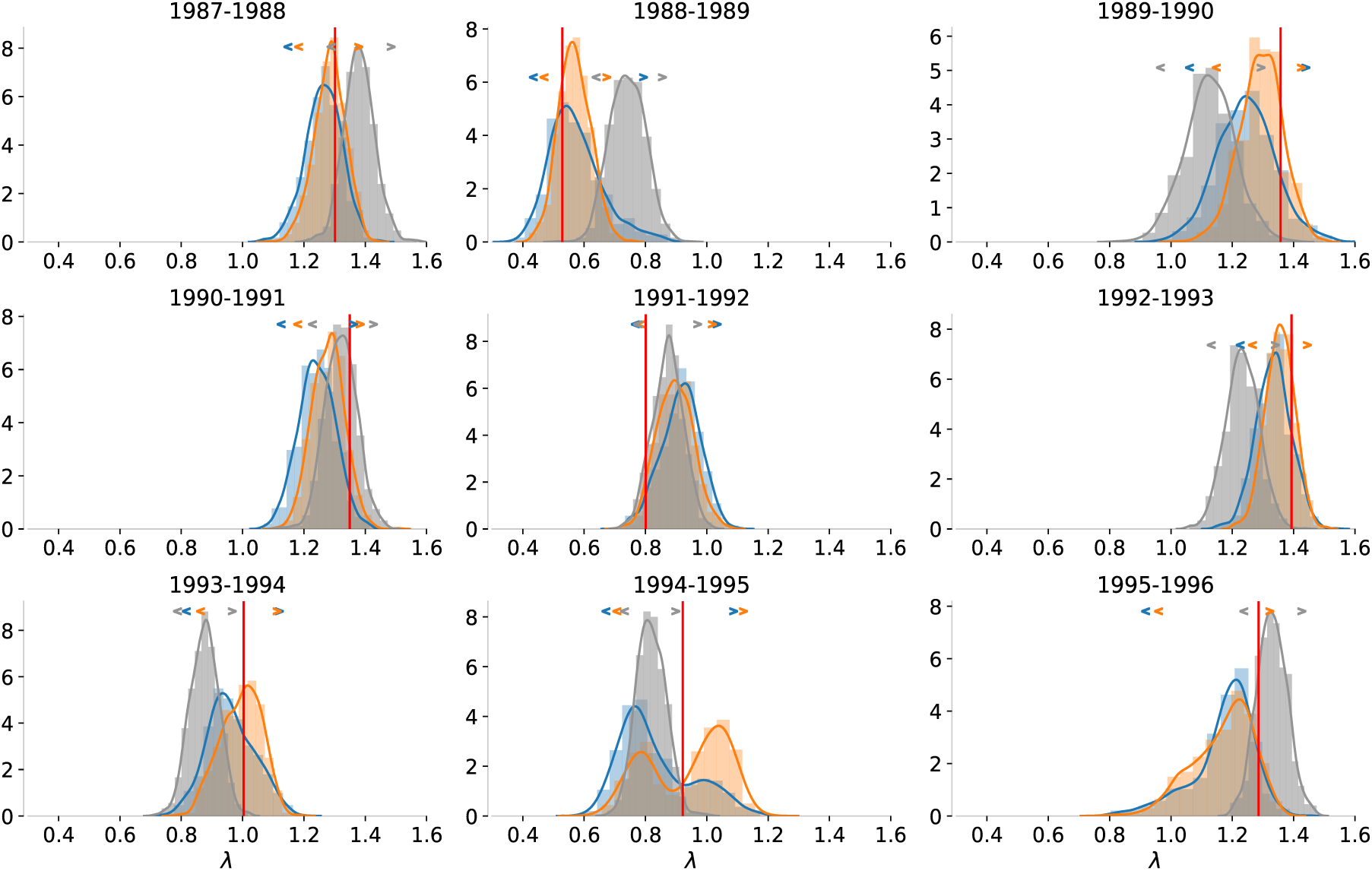
Histograms represent the density of simulated population growth rates generated through **1-step** forecasting. The associated 95% prediction intervals are denoted by angle brackets, each in its corresponding colour, positioned at the top of each histogram. The red vertical line represents the true observed population growth rate.

Consistent with earlier observations of *C. Flava*, GLM’s narrower intervals likely reflect a more constrained or rigid model structure. This does not necessarily translate to better accuracy. In fact, GLM often fails to include the true population growth rate within its P.I., particularly during the years 1988-1989-1990 and 1992-1993-1994-1995, revealing its limitations in capturing the underlying population dynamics. On the other hand, while GP and ABC_GP exhibit wider P.I., they are generally better at covering the true population growth rate, reflecting a better balance between uncertainty and accuracy.

A notable feature of ABC_GP is the bimodal distribution in 1994-1995, where two distinct peaks reflect different possible growth patterns contradicting to each other — one indicating population growth and the other suggesting shrinkage. The true population growth rate lies between these peaks, making both the mean and median predictions from ABC GP rather good for this year. This bimodal result may suggest that, constrained by the current ABC summary statistics, multiple plausible outcomes exist. It may also indicate a limitation of the proposed ABC sampler, as the fixed MCMC sample could prevent ABC from exploring a broader parameter space — potentially leading to fewer results concentrated between the peaks, where the true value is located (discussed in Zhu (2025).

## 6 Case study: Discussion and Future works

In this case study, building on previous research, we introduced novel covariates by modifying the NAO index and population density to reflect environmental factors affecting population dynamics. While these factors have been considered in past studies, our modifications represent a new contribution. These enhancements helped models to capture the general population size trends in line with the true observations. Additionally, incorporating ABC and GP methods provided more accurate point estimates and more reasonable prediction intervals, further improving performance compared to traditional approaches like GLMs.

However, during the ABC modeling, one key assumption was that the observed data for individual sheep each year were representative enough of the unmeasured population segments. Given the complexities observed in the capture process — such as the discrepancy between observed mothers and missing newborns (Section 3) — future research should incorporate domain-specific knowledge to propose a more appropriate and biologically meaningful capturing process. This could help bridge the gap between the simulated and observed datasets, improving their comparability during ABC sampling.

Additionally, to simplify this example as a demonstration of the `Python` package, we did not include age as a state variable, despite previous studies highlighting its noted impact on Soay sheep dynamics (Coulson et al. (2001); King et al. (2006); Ravindran et al. (2022)). Incorporating age in future models may hold potential of improving the accuracy of the predictions.

## 7 Discussion

This `Python` package provides a streamlined and user-friendly interface for implementing the ABC GP IPM approach. By integrating the selection of ABC summary statistics and the ABC sampling process, it simplifies complex workflows into several lines of `Python` code, making the method more accessible and allowing researchers to focus on their biological inquiries.

One of the key benefits of the package is its flexibility, allowing users to customise functions, such as those for IBMs. This adaptability allows it to accommodate specie-specific life cycles, making it suitable for various ecological contexts. However, users must follow specific conventions for inputs and outputs, promoting flexibility, reliability, and reproducibility.

While the package supports the early stages of demographic modeling, it does not include tools for constructing IPMs, as established R packages like `IPMpack` and `ipmr` have addressed this need. For those integrating `Python` with `R`, tools like `rpy2` can facilitate inter-operability. However, a primary limitation is the use of `GPflow` for GP modeling, which is deeply integrated with `TensorFlow` in `Python`. Currently, there is no straightforward way to transfer fitted GP models directly to `R`, meaning predictions and outputs must be completed in `Python` before transferring results to `R` for further analysis.

Additionally, most established IPM tools in `R` are designed to accept direct model structures, such as formulae describing GLMs, and are not equipped to incorporate predictions from external sources, such as predictions about meshpoints generated by `Python`-based models. While the online guide of our package offers `Python` scripts to assist in constructing IPM matrices from these predictions, this solution requires researchers to either adapt their existing `R` workflows to include `Python` components or manage separate workflows in both programming languages. Users who primarily work with `R` may find the approach challenging.

## Code availability

The package can be installed via pip or from PyPI at https://pypi.org/project/abc-gp-ipm/0.0.1/.

## Acknowledgements

For the purpose of Open Access, the author has applied a CC BY public copyright licence to any Author Accepted Manuscript version arising from this submission.

## A Appendix

For the case study of Soay sheep, each vital rate was modelled separately as a function of the individual’s state in the current year (*z*) or the following year (*z^′^*), along with environmental covariates ***ω*** representing modified weather and population density factors. Specifically:

- probabilities of survival *s*(*z|****ω***) and probabilities of recruitment *p_f_* (*z|****ω***), using GPs with Bernoulli likelihoods;
- growth function *G*(*z^′^|z,* ***ω***) and newborn size distribution *r_size_*(*z^′^|z,* ***ω***), using GP regressions.

For each of the GP models, the mean function was set to zero, and the Gaussian kernel function was applied. The Gaussian kernel function was employed with the automatic relevance determination setting, allowing it to separately modulate the magnitude of similarity in each input coordinate (size, modified weather and density factors). Bayesian individual vital rate model fitting was conducted using non-informative priors and Hamiltonian Monte Carlo methods (Hensman et al. (2015)). All the GP model fitting tasks were implemented using the `GPflow` package in the `Python` programming language (Van Rossum & Drake Jr (1995); Matthews et al. (2017)).

**Figure 5:**
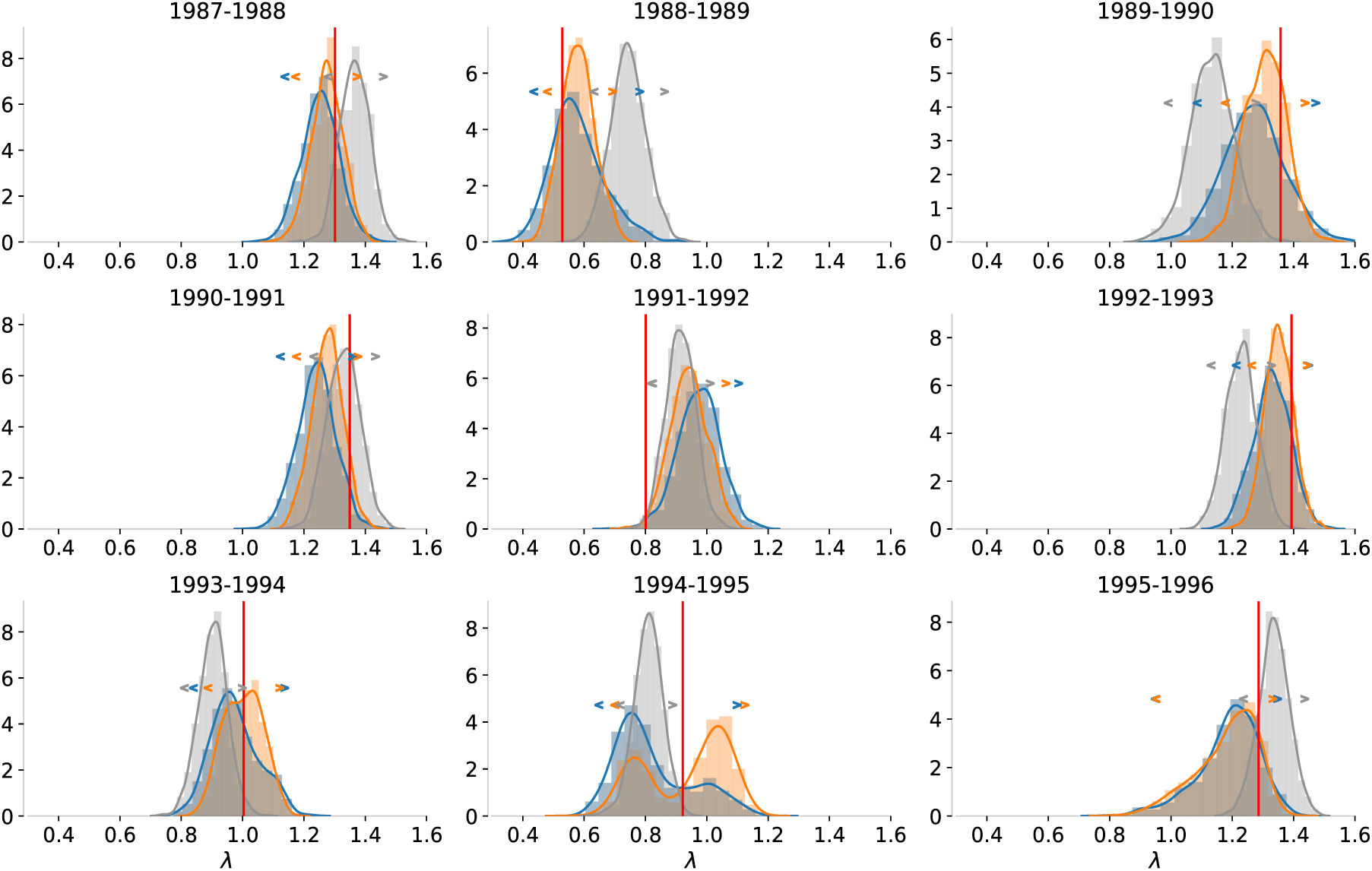
Histograms represent the density of simulated population growth rates generated through **Long-term** forecasting. The associated 95% prediction intervals are denoted by angle brackets, each in its corresponding colour, positioned at the top of each histogram. The red vertical line represents the true observed population growth rate.

